# Transcriptionally defined AML cell states associate with treatment response and microenvironmental remodeling

**DOI:** 10.64898/2026.07.01.735780

**Authors:** Nona Struyf, Leonard Hartmanis, Lucia Rico Pizarro, Albin Österroos, Anna Bohlin, Sofia Bengtzén, Sören Lehmann, Olli Kallioniemi, Tom Erkers

## Abstract

While therapy resistance in acute myeloid leukemia (AML) is often attributed to leukemic stem cells (LSCs), their functional properties are not fully captured by their well-established genetic landscape and cell lineage transcriptional programs. Here, we explore AML cell states and their associations to drug response and systemic immune context.

We performed integrated single-cell transcriptomics and immunophenotyping on diagnostic AML samples (n=6) to define transcriptional cell state gene signatures. These were projected onto bulk RNA-seq data from 448 AML patients to assess associations with drug sensitivity, plasma proteomics, clinical features, and established prognostic scores. Longitudinal single-cell data from external cohorts and cell-cell communication analyses were used to examine treatment dynamics and microenvironmental signaling.

We defined nine AML cell states, including progenitor-like, stromal-like, antigen-presenting, and monocytic programs. Stemness features were distributed across multiple states, with lymphoid-primed and stress-adapted progenitors showing the strongest alignment with established stemness scores. Distinct drug sensitivities emerged, including cell cycle checkpoint inhibitor sensitivity in stress-adapted progenitors and kinase inhibitor sensitivity in cycling progenitors, alongside shared resistance to BH3 mimetics in monocytic states. Stress-adapted progenitors were associated with adverse clinical features and expanded following venetoclax-based therapy. Monocytic states acted as immunosuppressive hubs via TIGIT signaling, while stromal-associated states received niche-derived survival signals.

Overall, we define a framework that associates AML cell states with stemness, drug response, and microenvironmental interactions. These findings highlight distributed stemness, state-specific vulnerabilities, and niche-driven resistance mechanisms, informing more precise therapeutic strategies in AML.

## Introduction

Acute myeloid leukemia (AML) is the most common form of acute leukemia in adults, affecting myeloid progenitor cells in blood and bone marrow.^1^ Although AML is treated with intensive induction chemotherapy consisting of daunorubicin and cytarabine, relapse rates remain high. Older patients unfit for intensive chemotherapy are usually given palliative or supportive care consisting of hypomethylating agents (HMA) with or without venetoclax, or low-dose cytarabine (LDAC).^2^

The main driver of therapy resistance in AML are leukemic stem cells (LSCs) which are dysfunctional hematopoietic stem cells (HSPCs) that have the ability for self-renewal and differentiation. LSCs have been classically defined as CD34^+^CD38^-^ cells that can initiate leukemia after transplantation into immunodeficient mice.^3,4^ However, this phenotype is not consistently expressed across patients.^5,6^ Additionally, stem-like behavior has been found in cells outside of this classical immunophenotype, and transcriptional programs are not able to map cleanly onto surface marker-defined populations.^7^ Other surface markers such as CD200 and CD45RA have also been shown to be present on LSCs, indicating that important LSC subsets may be overlooked.^8–10^ There is additional evidence that within each AML patient there may be multiple functionally heterogeneous LSC clones present.^11,12^ This suggests that the functional properties attributed to LSCs may be better understood as features of broader transcriptional cell states rather than properties confined to a distinct immunophenotypic population.

LSC survival and proliferation are supported by the bone marrow (BM) microenvironment, where interactions with mesenchymal stromal cells, endothelial cells, and immune populations create a protective niche that promotes quiescence and therapy resistance.^13^ Beyond providing passive support, leukemic cells actively remodel the BM niche through dysregulated cytokine and growth factor signaling, creating feedback loops that further sustain LSC survival and suppress normal hematopoiesis.^14^ Additionally, specific genetic mutations have been shown to alter cytokine signaling between leukemic cells and the tumor microenvironment, potentially causing therapy resistance even in initially responsive patients.^15,16^ Importantly, cell-cell communication between leukemic populations and their niche can vary. Recent advances in single-cell RNA sequencing (scRNA-seq) have begun to reveal the transcriptional complexity of AML at unprecedented resolution. Studies have identified distinct leukemic cell populations within individual patients that mirror normal hematopoietic hierarchies but display aberrant transcriptional programs, suggesting that AML is better understood as a collection of coexisting cellular states rather than a single malignancy.^17–20^ These approaches have further demonstrated that transcriptionally defined states capture functional heterogeneity that bulk RNA sequencing cannot resolve, and that such states may have distinct clinical implications. However, a comprehensive functional classification of AML cell states that integrates transcriptional identity with microenvironmental signaling, drug response, and clinical outcome has not yet been established.

In this study, we define distinct functional states of AML cells and investigate how they may modulate their surrounding microenvironment. Additionally, we show how these states relate to therapy response and clinical outcomes in AML. To this end, we utilized scRNA-seq combined with CITE-seq to define transcriptional AML states and generate robust transcriptomic signatures. We then utilized a bulk RNA-seq data comprising of 448 AML patients, whereby applying these signatures we could link AML cell state-specific profiles to ex vivo drug response, clinical outcomes, and soluble protein levels. Similarly, using publicly available scRNA-seq datasets, we validated the identified AML cell states and explored their functional dynamics before and after treatment. Finally, we demonstrate the potential microenvironmental influence these cell states may have on in an autologous and paracrine manner, which may potentiate drug resistance mechanisms.

## Results

### Single-cell transcriptomes define AML programs with supporting immunophenotypes

The initial dataset consisted of scRNAseq and immunophenotyping on bone marrow obtained from six AML patients at diagnosis (Figure 1A). First, we performed cell type annotation using the BoneMarrowMap^21^ (BMM) reference atlas. Cells labeled as pure lymphocyte populations according to the BMM were removed (Supplementary figure 1A). This was followed by unsupervised clustering of the myeloid cells, which resulted in 16 primary clusters that largely capture the inter-patient heterogeneity as is expected in AML (Figure 1B, top, Supplementary figure 1B). Using the BMM, we could then map the predicted cell type for each single cell and the dominant cell type per cluster, showing clustering of monocytic and precursor cell subsets (Figure 1B, bottom, Supplementary figure 1C). This resulted in five main cell type annotations: Cycling progenitors, Early lymphoid cells, HSC multipotent progenitors (MPPs), Monocytes, and Stromal cells. Next, we scored each individual cell for known HSC/LSC biology signatures.^17,18^ All but the monocyte clusters scored higher for quiescent LSPC, primed LSPC, and HSC signatures (Figure 1C). The HSC MPP, Stromal, and Cycling progenitor assigned clusters scored the highest for cycle LSPCs, but more variation could be observed in the Monocyte populations, indicating heterogeneity and potential monocyte sub-populations. Even though the cells occupy patient-specific manifolds as expected due to the large heterogeneity in AML, we can observe that they express conserved transcriptional programs leading to similar cell type assignments.

**Figure 1:**
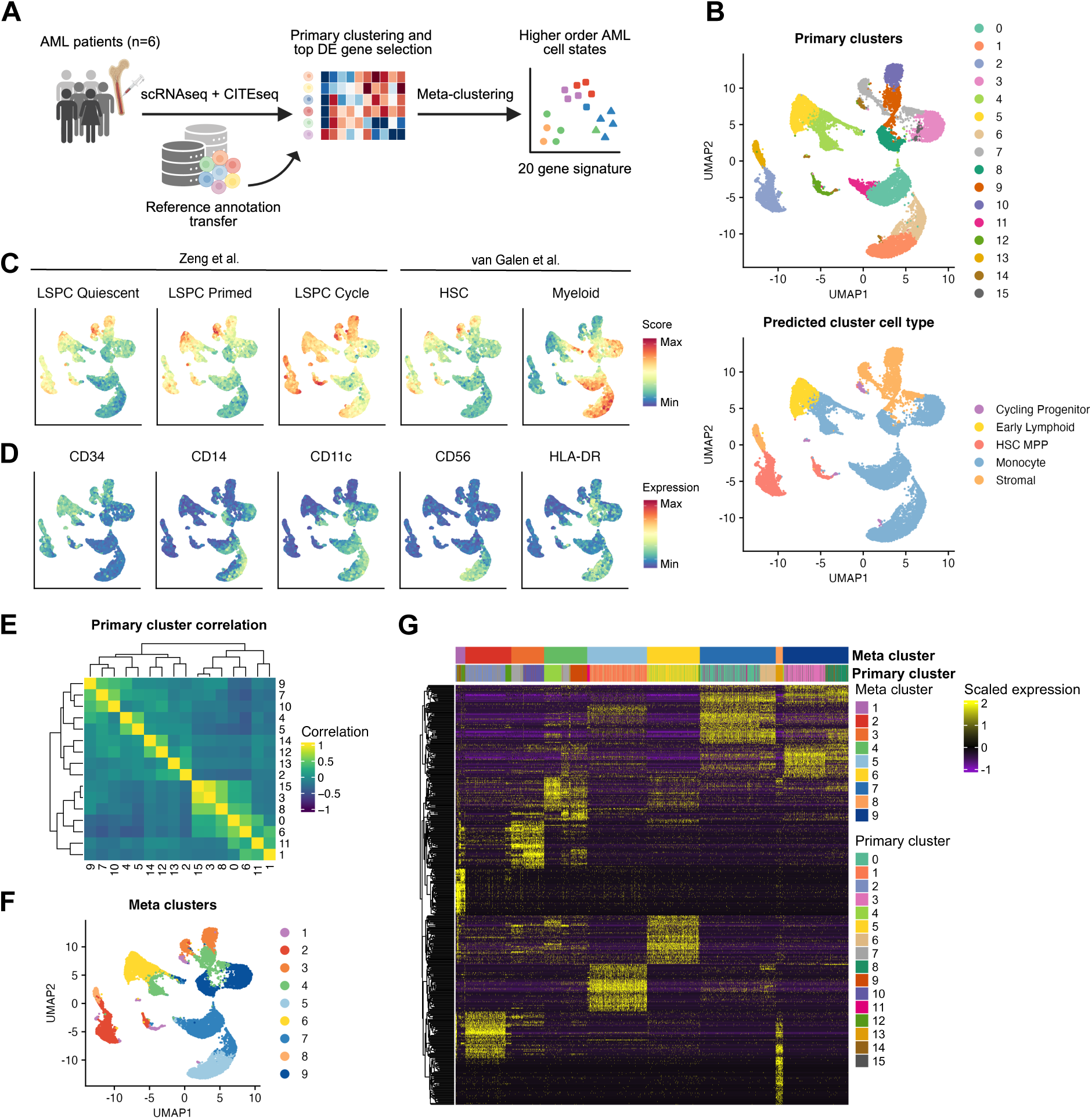
Integration of AML patient scRNAseq and CITE-seq data with cell type atlases to define transcriptional meta-clusters supported by immunophenotyping. (A) Schematic overview of data integration and clustering. (B) UMAP of all myeloid cells from AML patients (n=6). Cells are colored by unsupervised clusters obtained by graph-based clustering (Louvain alghorithm) in Seurat using the first 20 principal components (top) or broad predicted cell type based on the BoneMarrowMap atlas (bottom). (C) UMAP overlaid with enrichment scores for established AML HSC/LSC signatures. (D) UMAP overlaid with surface protein expression measured by CITE-seq. (E) Pearson correlation matrix showing correlation between primary clusters based on the top 50 DE genes per cluster. (F) UMAP of single cells colored by meta-clusters determined from K-means (k=9) clustering of the top principal components of top 50 DE genes per cluster. (G) Heatmap showing scaled expression of top 50 DE genes per meta-cluster showing correspondence between meta– and primary clusters.

Furthermore, we could observe clear concordance between cell surface marker expression and transcriptomics. The Cycling progenitor, Early lymphoid, HSC MPP, and Stromal clusters expressed higher levels CD34 and CD117 while the Monocytic clusters expressed CD11c, CD14, and CD15, albeit at different intensities (Figure 1D, Supplementary Figure 1D). Interestingly, the monocyte heterogeneity was also captured by some degree of CD34 expression, as well as expression of the NK-cell marker CD56 on subgroups, a known prognostic indicator in AML.^22^ Similarly, varying levels of HLA-DR are expressed in both the Monocyte clusters, as well as in subsets of the Stromal clusters. These results indicate both common biology between the primary clusters, as well as immunophenotypic subpopulations within similarly assigned cell types.

To assess transcriptional similarity between the primary clusters, we calculated pairwise Pearson correlations using the top 50 differentially expressed genes for each cluster. By restricting the analysis to this selection of cluster-defining genes, shared biological programs are emphasized rather than global similarity driven by housekeeping genes or patient specific expression. This analysis revealed clusters with higher transcriptional similarity, motivating the aggregation of clusters into higher-order transcriptional states (Figure 1E). We then performed meta-clustering of primary clusters using the cluster-defining genes. The optimal number of meta-clusters was determined by silhouette width analysis, resulting in the selection of nine transcriptionally coherent meta-clusters (Figure 1F, Supplementary figure 1E). Next, we identified the top differentially expressed genes for each meta-cluster which revealed coherent gene expression patterns within meta-clusters, while also highlighting their composition from multiple primary clusters (Figure 1G).

### AML cell states display distinct functional identities and stemness distributions

To explore the biological identity of the nine established meta-clusters, we examined the expression of representative marker genes derived from the most differentially expressed genes within each cluster. While several clusters shared lineage-associated genes such as CD34, CD38, and CD14, each was characterized by distinct enrichment of specific gene expression markers (Figure 2A-C, Supplementary Figure 2A). We hereon refer to the clusters as cell states, where each cell state was assigned a name based on BMM assigned cell lineage, key gene/protein markers, and overrepresented pathways. In order by cluster; Cycling progenitor (C-Prog), Stress-adapted progenitor (S-Prog), Vascular-like stromal (VLS), Antigen-presenting myeloid (APM), Cytotoxic effector monocyte (CEM), Lymphoid-primed progenitor (LPP), Cytokine-responsive monocyte (CRM), Mesenchymal-like stromal (MLS), and Tissue-remodeling inflammatory monocyte (TIM) (Figure 2D).

**Figure 2:**
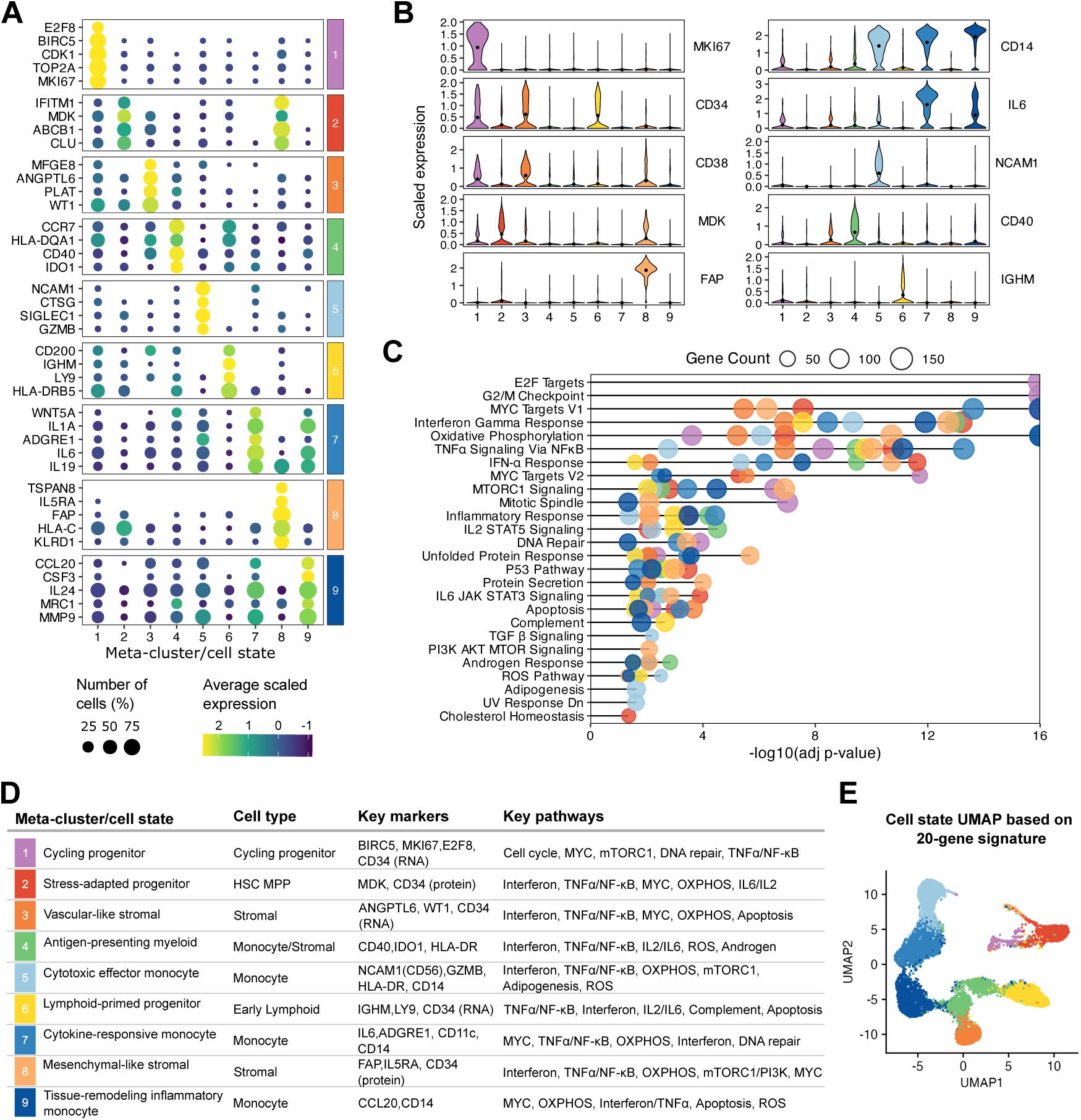
Cells of each meta-cluster display broad cell type association but distinct functional specializations. (A) Dotplot showing a selection of top DE genes for each meta-cluster colored by average scaled expression while dot size represents the percentage of cells where each marker is expressed. (B) Violin plots of selected genes shown as scaled expression. (C) Dot plot showing enriched pathways in each cell state determined by over-representation analysis (ORA) performed with the Hallmark gene sets from MSigDB. Pathways with significant enrichment (adjusted p < 0.05) are shown with dot size corresponding to the number of genes in the pathway, and color corresponding to cell state identity. (D) Summary table of each cell state with description, assigned BMM cell type, key markers determined by scRNAseq and CITE-seq, and key enriched pathways. (E) UMAP showing single cells mapped by 20-gene signature.

C-Prog cells exhibited signs of a proliferative AML cell state defined by expression of cell cycle and mitotic regulators, including MKI67, TOP2A, CDK1, BIRC5, and multiple kinesins and checkpoint-associated genes indicating a proliferation-driven state. This transcriptional program is consistent with actively cycling progenitor-like leukemic cells and expressed both CD34 and CD38. S-Prog cells expressed interferon– and survival-related genes such as IFITM1, MDK, ABCB1, and CLU. Even though we observed surface protein expression of CD34 in this cell state, RNA levels were lower compared to other progenitor-like states. The LPP cell state was similarly assigned as early lymphoid progenitor transcriptional phenotype. Although marked by gene expression of CD34, it also showed expression of IGHM, LY9, HLA-DRB5, and CD200.

The two stromal-associated clusters exhibited partially overlapping transcriptional profiles. However, these states remained distinguishable by cluster-specific marker expression. VLS cells were characterized by expression of vascular-like stromal associated genes such as PLAT, ANGPTL6, and MFGE8, whereas MLS cells showed enrichment of mesenchymal associated genes such as IL5RA, FAP, and LMOD1. Additionally, VLS showed high CD34 and CD38 gene expression, while MLS had low CD34 but intermediate CD38 expression.

Several cell states displayed programs consistent with monocytic differentiation. CEM was defined by CD56 (NCAM1) protein and gene expression, along with high expression of GZMB and CTSG. CRM was characterized by enrichment of cytokine– and chemokine-associated genes, including IL6, IL19, WNT5A, and FPR2. In contrast, TIM showed preferential expression of genes such as MRC1, MMP9, CCL20, and FGF2, found in niche-interacting or macrophage-like phenotypes. All three displayed CD14 expression at different levels, with TIM showing high expression while CEM had the lowest.

Notably, the APM cell state exhibited features of both monocytic and stromal programs, with expression of antigen presentation and immune regulatory genes including CD40, CCR7, IDO1, and multiple HLA-DQ family members. Moreover, APM was the only mature myeloid cell state without CD14 surface protein expression but high protein expression of HLA-DR (Figure 1D).

A limited subset of these markers was sufficient to distinguish each AML cell state, motivating the derivation of a reduced 20-gene signature that preserved state-specific transcriptional identity across datasets (Figure 2E, Supplementary table 1). These signatures show that while certain patients are more dominant for certain cell states, the cell states are not patient specific (Supplementary Figure 2B).

To assess the specificity of the gene signatures, we compared their classification performance to randomly generated gene sets of matched size. The true signatures achieved high accuracy in recapitulating cluster identities (81%), whereas random gene sets performed at near-chance levels (13%, Supplementary Figure 2C-F). Misclassification primarily occurred between transcriptionally related meta-clusters such as C-Prog and S-Prog.

### AML cell state enrichment associates with clinical outcome

To link the AML cell state-specific profiles with clinical outcomes, ex vivo drug response, and soluble protein expression in a larger cohort, we utilized a bulk RNA-seq data comprising of 448 AML patients (Figure 3A). We used the 20-gene cell state signatures and performed GSVA on all samples which resulted in a matrix with per-patient enrichment score for each cell state (Figure 3B). We then examined associations between GSVA-derived cell state scores and clinical parameters across all patients with available clinical annotation. Spearman correlation analysis of continuous variables revealed that LPP (ρ=-0.46, p_adj=1.1×10^-^⁸), APM (ρ=-0.39, p_adj=2.8×10^-6^), S-Prog (ρ=-0.35, p_adj=3.1×10^-5^), C-Prog (ρ=-0.31, p_adj=2.5×10^-4^), and VLS (ρ=-0.29, p_adj=7.2×10^-4^) enrichment were most strongly and inversely associated with white blood cell (WBC) count, while CRM (ρ=-0.47, p_adj=1.1×10^-^⁸), TIM (ρ=-0.36, p_adj=2.7×10^-5^), CEM (ρ=-0.33, p_adj=9.75×10^-5^), and APM (ρ=-0.32, p_adj=2.1×10^-4^) enrichment were inversely correlated with bone marrow blast percentage (Figure 3C, Supplementary table 2). This suggests broad depletion of progenitor, immune, and stromal-like states in high disease burden AML. S-Prog enrichment showed a weak positive correlation with age (ρ=0.28, p_adj=2.5×10^-4^), while MLS enrichment was negatively correlated with age (ρ=-0.19, p_adj=2.4×10^-2^).

**Figure 3:**
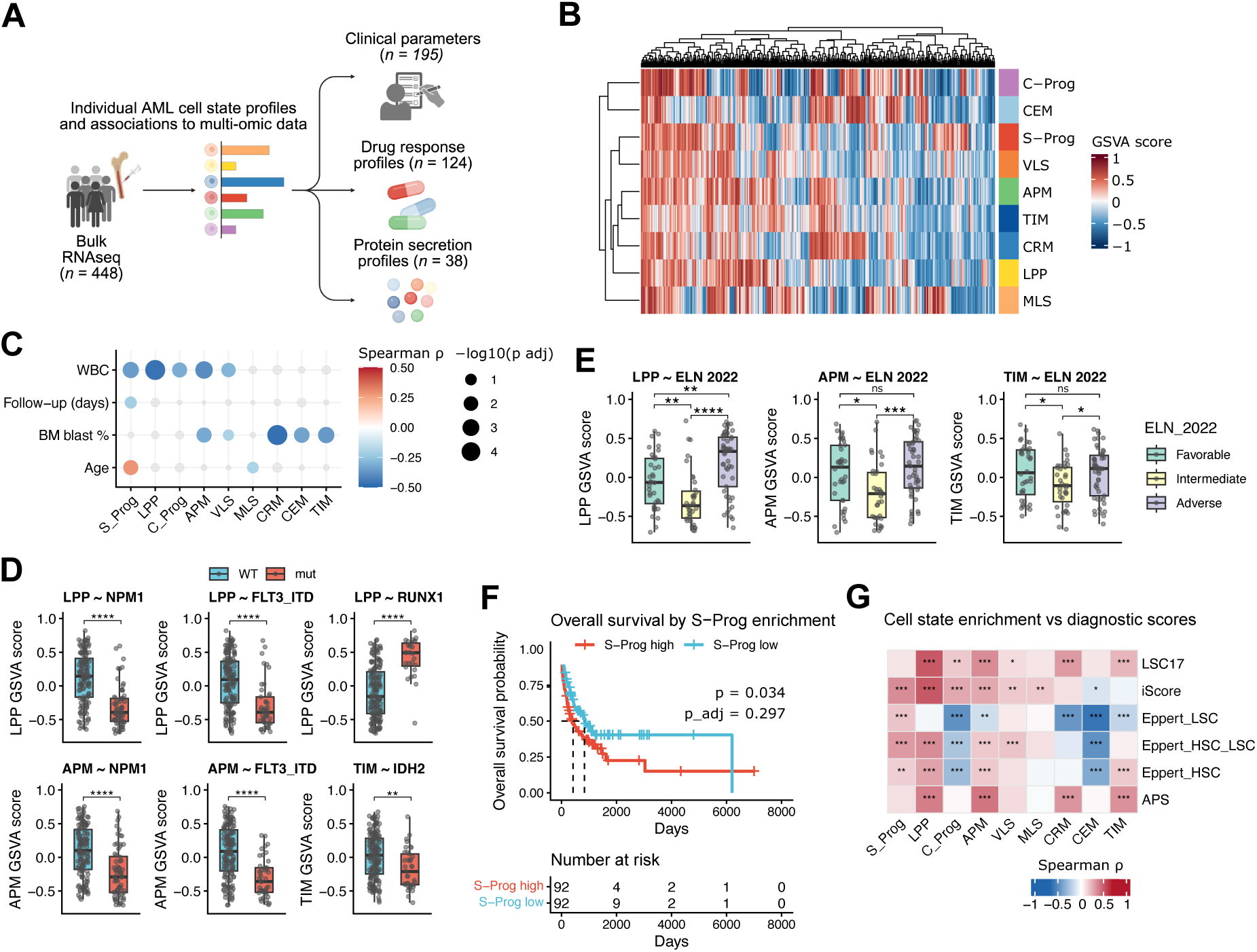
Clinical associations with the nine AML cell states. (A) Visual representation of the analysis workflow for application of cell state signatures to an independent cohort bulk RNAseq data. (B) Heatmap displaying the cell state GSVA scores per patient (n=448). (C) Associations of continuous clinical variables with significant correlations to the cell state GSVA scores. (D) Boxplots of mutational and (E) ELN 2022 status and associations to the cell state GSVA scores, significance was assessed with the Wilcoxon test. (F) Kaplan-Meier curve for S-Prog Cell state association and survival. (G) Heatmap with Spearman ρ values showing correlation between cell state GSVA scores and known LSC signatures.

Mutation-specific associations were identified across multiple states using Wilcoxon rank-sum testing (Figure 3D, Supplementary figure 3A, Supplementary table 3). LPP and APM showed the strongest and most consistent mutation associations, with both states significantly depleted in NPM1-mutant patients (LPP: p_adj=5.5×10^-11^; APM: p_adj=3.4×10^-6^) and FLT3-ITD-positive patients (LPP: p_adj=2.6×10^-6^; APM: p_adj=1.4×10^-7^). TIM and VLS were additionally depleted in FLT3-ITD-positive patients (TIM: p_adj=2.6×10^-4^; VLS: p_adj=1.6×10^-2^), and CRM showed a similar trend (p_adj=3.0×10^-2^). In contrast, LPP enrichment was significantly elevated in RUNX1-mutant AML (p_adj=1.6×10^-7^), consistent with lymphoid-primed transcriptional programs described in this subtype.^23^ IDH2 mutation was associated with depletion of both TIM (p_adj=2.0×10^-2^) and MLS (p_adj=2.2×10^-2^).

Kruskal-Wallis analysis across ELN 2022 risk groups revealed consistent enrichment of progenitor and immune states in Intermediate risk patients (Figure 3E, Supplementary table 4). LPP showed the strongest ELN association (Adverse vs Intermediate p_adj=1.1×10^-6^, Favorable vs Intermediate p_adj=1.9×10^-2^), followed by APM (Adverse vs Intermediate p_adj=7.1×10^-4^) and TIM (both comparisons p_adj < 0.05). C-Prog was similarly elevated in Intermediate compared to both other groups (Supplementary Figure 3C). Notably, S-Prog enrichment was significantly higher in Adverse compared to Favorable risk patients (p_adj=3.9×10^-2^).

Kaplan-Meier analysis using median GSVA enrichment as a dichotomization threshold identified S-Prog as the state most strongly associated with overall survival, with high S-Prog enrichment associated with significantly shorter survival in the unadjusted analysis (median OS: 299 vs 489 days, log-rank p=0.034, Figure 3F, Supplementary table 5). LPP and CEM showed trends in the same direction (p=0.082 and p=0.099), though neither reached significance (Supplementary figure 3B). No state survived correction for multiple testing across all nine states, likely reflecting limited statistical power given incomplete survival annotation available for 184 of 448 bulk cohort patients.

To contextualize AML cell state enrichment relative to established diagnostic or HSC/LSC scoring frameworks, we calculated the LSC17^24^, iScore^19^, and APS^25^ as well as the classical Eppert LSC/HSC^7^ signatures for all 448 bulk RNA-seq patients and examined Spearman correlations between these scores and state-specific GSVA enrichment (Figure 3G, Supplementary Figure 3D). LPP and S-Prog showed significant positive associations across almost all five diagnostic scores, with LPP demonstrating the strongest correlation with iScore (ρ=0.50, p_adj<0.001) and LSC17 (ρ=0.49, p_adj<0.001), and S-Prog correlating with iScore (ρ=0.34, p_adj<0.001), reinforcing their stem-like transcriptional identity across independent frameworks. CEM showed several negative associations, significantly anti-correlating with Eppert HSC (ρ=-0.39, p_adj<0.001), Eppert LSC (ρ=-0.62, p_adj<0.001) and Eppert HSC-LSC (ρ=-0.49, p_adj<0.001) signatures, indicating that CEM-enriched patients have uniformly low classical stemness scores. C-Prog showed negative correlations to Eppert LSC (ρ=-0.50, p_adj<0.001) and Eppert HSC (ρ=-0.31, p_adj<0.001) signatures while positively associating with iScore (ρ=0.26, p_adj<0.001). The associations of each cell state with the more updated APS followed those of LSC17 closely, however, APM showed a higher positive association (APS: ρ=0.42, p_adj<0.001, LSC17: ρ=0.34, p_adj<0.001). This data demonstrates that transcriptional cell state enrichment captures clinically relevant heterogeneity that is partially but incompletely represented by existing diagnostic scores.

### Transcriptional AML cell states associate with distinct drug sensitivity profiles and plasma protein signatures

To assess the functional relevance of the nine AML cell states, we associated the per-patient cell state enrichment score with ex vivo drug testing data (n=124) and soluble proteomics (n=38). We calculated Spearman correlations between the enrichment scores and selective drug sensitivity scores (sDSS, Figure 4A-B, Supplementary table 6). Unsupervised hierarchical clustering of the resulting correlation matrix revealed that drug sensitivity profiles grouped broadly by lineage identity, with progenitor states (S-Prog, LPP, APM, C-Prog), stromal states (MLS, VLS), and monocytic states (CRM, CEM, TIM) forming partially distinct clusters. Although APM consists of a mixed stromal/monocytic phenotype, its drug response profile more closely resembled the progenitor-like states. Additionally, the monocytic states showed a more resistant profile with some sensitivity to MEK-inhibitors (e.g. selumetinib, trametinib), an association that has been previously described in monocytic AML. S-prog enrichment was positively associated with sensitivity to the Wee1 inhibitor AZD1775 (ρ=0.43, p_adj=0.021) and the PLK1 inhibitor volasertib (ρ=0.33, p_adj=0.041), and was negatively associated with the BET inhibitor dBET1 (ρ=-0.47, p_adj=0.015, Figure C). C-Prog enrichment was associated with overall drug resistance, consistent with prior reports linking cycling progenitor states to poor prognosis in AML.^18^ However, C-Prog enrichment was significantly associated to crenolanib sensitivity (ρ=0.32, p_adj=0.18), a FLT3 tyrosine kinase inhibitor. This increase in sDSS was similar in both FLT3-ITD+ and WT groups, suggesting C-Prog enrichment identifies crenolanib-sensitive disease independently of FLT3 mutation status (Supplementary figure 4A).

**Figure 4:**
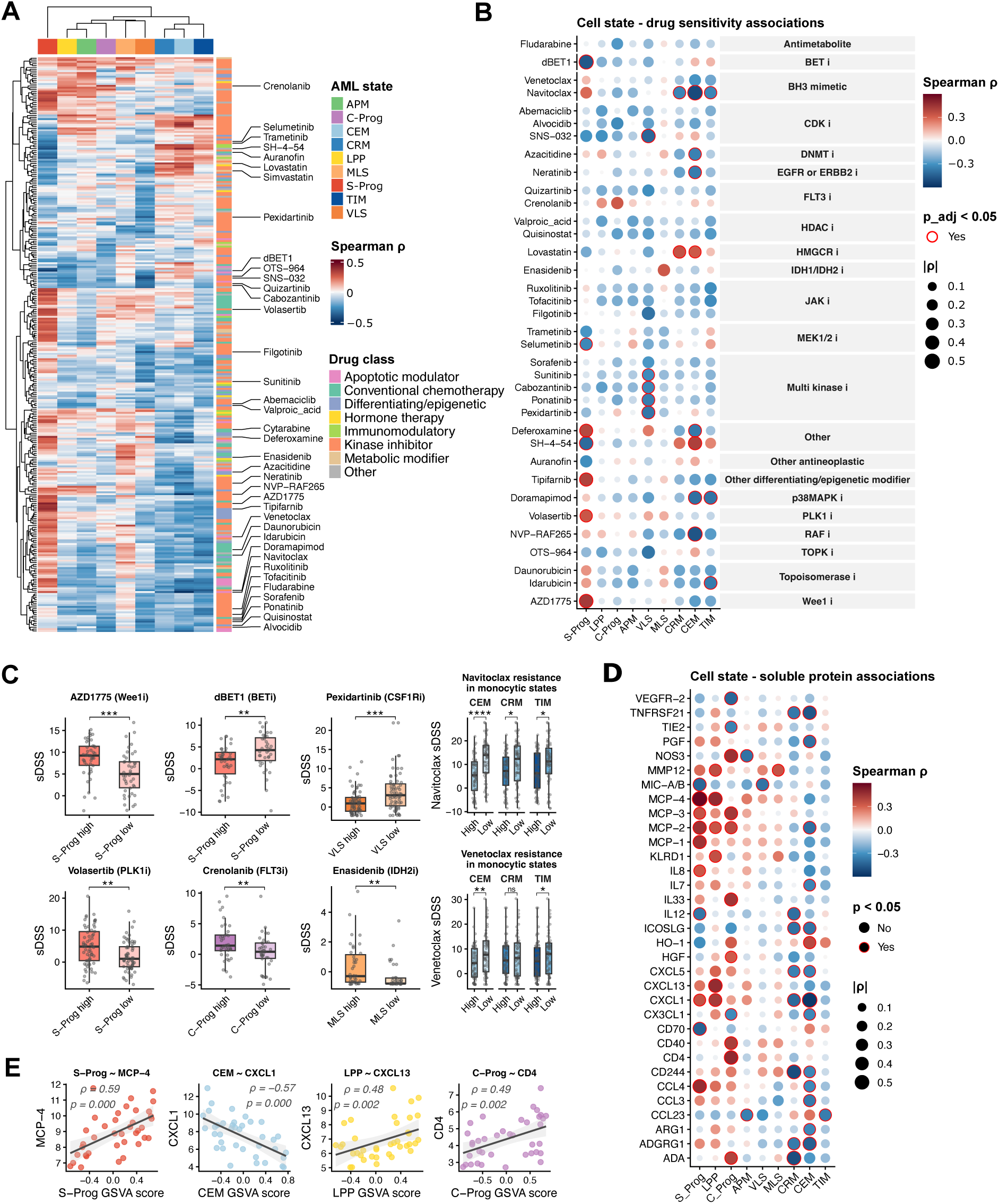
Cell states are associated with ex vivo drug response and plasma protein profiles. (A) Heatmap of drugs significantly (p<0.05) associated to cell state GSVA scores displayed as Spearman ρ. (B) Dotplot of selected drugs with cell state GSVA associations shown as Spearman ρ, with size indicating absolute ρ, color showing ρ directionality, and external red border showing associations with adjusted p<0.05, determined by Benjamini-Hochberg procedure. (C) Boxplots of selected drug-cell state associations, significance was assessed with the Wilcoxon test. (D) Dotplot of selected soluble proteins with cell state GSVA associations shown as Spearman ρ, with size indicating absolute ρ, color showing ρ directionality, and external red border showing associations where p<0.05, determined by Spearman rank correlation. (E) Scatterplots showing association between selected soluble proteins and cell state GSVA scores for all tested patients (n=38), assessed with Spearman correlation.

Out of the stromal states, MLS enrichment showed a trend toward resistance to the IDH2 inhibitor enasidenib (ρ=0.33, p_adj=0.017), consistent with the significant depletion of MLS in IDH2-mutant patients observed in our clinical association analysis. We could also confirm that enasidenib sensitivity was almost exclusive to IDH2 WT patients (Supplementary figure 4B). VLS on the other hand, showed resistance to multiple kinase inhibitors including pexidartinib (CSF1R inhibitor, ρ=-0.28, p_adj=0.015), ponatinib (ρ=-0.36, p_adj=0.021), sunitinib (ρ=-0.32, p_adj=0.047), and cabozantinib (ρ=-0.33, p_adj=0.041) (Supplementary figure 4C).

Within the monocytic clusters, resistance to the BH3 mimetic Navitoclax was the most consistent and statistically robust finding, with the strongest association observed in CEM (ρ=-0.51, p_adj<0.0001) followed by TIM (ρ=-0.34, p_adj=0.041) and CRM (ρ=-0.34, p_adj=0.040). Despite partial inverse co-enrichment between CEM and CRM at the patient level and independent enrichment of TIM, all three monocytic states converged on BH3 mimetic resistance. A similar trend was observed for Venetoclax across monocytic states although no significance could be observed (CEM: ρ=-0.29, p_adj=0.080, TIM: ρ=-0.24, p_adj=0.216, CRM: ρ=-0.15, p_adj=0.511).

Several states showed significant soluble protein associations that were consistent with their transcriptional identity and drug sensitivity profiles (Figure 4D-E, Supplementary table 7). S-Prog enrichment showed the strongest overall association with MCP-4 (ρ=0.59, p=0.0001) alongside positive associations with MCP-1, MCP-2, MCP-3, CCL4, and CXCL1, indicating that S-Prog enriched patients have elevated monocyte-recruiting chemokine levels. Conversely, S-Prog enrichment was negatively associated with cytotoxic immune activation markers CD70, MIC-A/B, and IL-12.

CEM enrichment showed the richest protein signature, with CXCL1 representing the strongest negative association (ρ=−0.573, p=0.0002), alongside depletion of immune activation markers including ADGRG1, TNFRSF21, ICOSLG, CCL3, CCL4, and CXCL5. The single positive CEM association was HO-1 (ρ=0.38, p=0.020), an immunosuppressive enzyme involved in heme catabolism and immune tolerance.

LPP enrichment was positively associated with CXCL13 (ρ=0.48, p=0.003), a B cell-recruiting chemokine, alongside MCP-4, CXCL1, MMP12, and KLRD1. The CXCL13 association is particularly coherent with LPP’s lymphoid-primed transcriptional identity. C-Prog enrichment associated with elevated CD4 (ρ=0.49, p=0.002), NOS3, CD40, and IL-33 alongside reduced TIE2 and VEGFR-2. Stromal states VLS and MLS showed more limited protein associations, both showed weak positive associations to MMP12 (VLS: ρ=0.23, p=0.168, MLS: ρ=0.35, p=0.034), although no significance was reached.

### Functional AML cell states display treatment-specific compositional dynamics

We next sought to explore how these states respond to treatment at single-cell resolution. Using two publicly available scRNA-seq datasets, we applied our 20-gene state signatures to individual cells using AUCell scoring and assigned each cell to its most dominant state, enabling direct tracking of state composition before and after therapy (Figure 5A).

**Figure 5:**
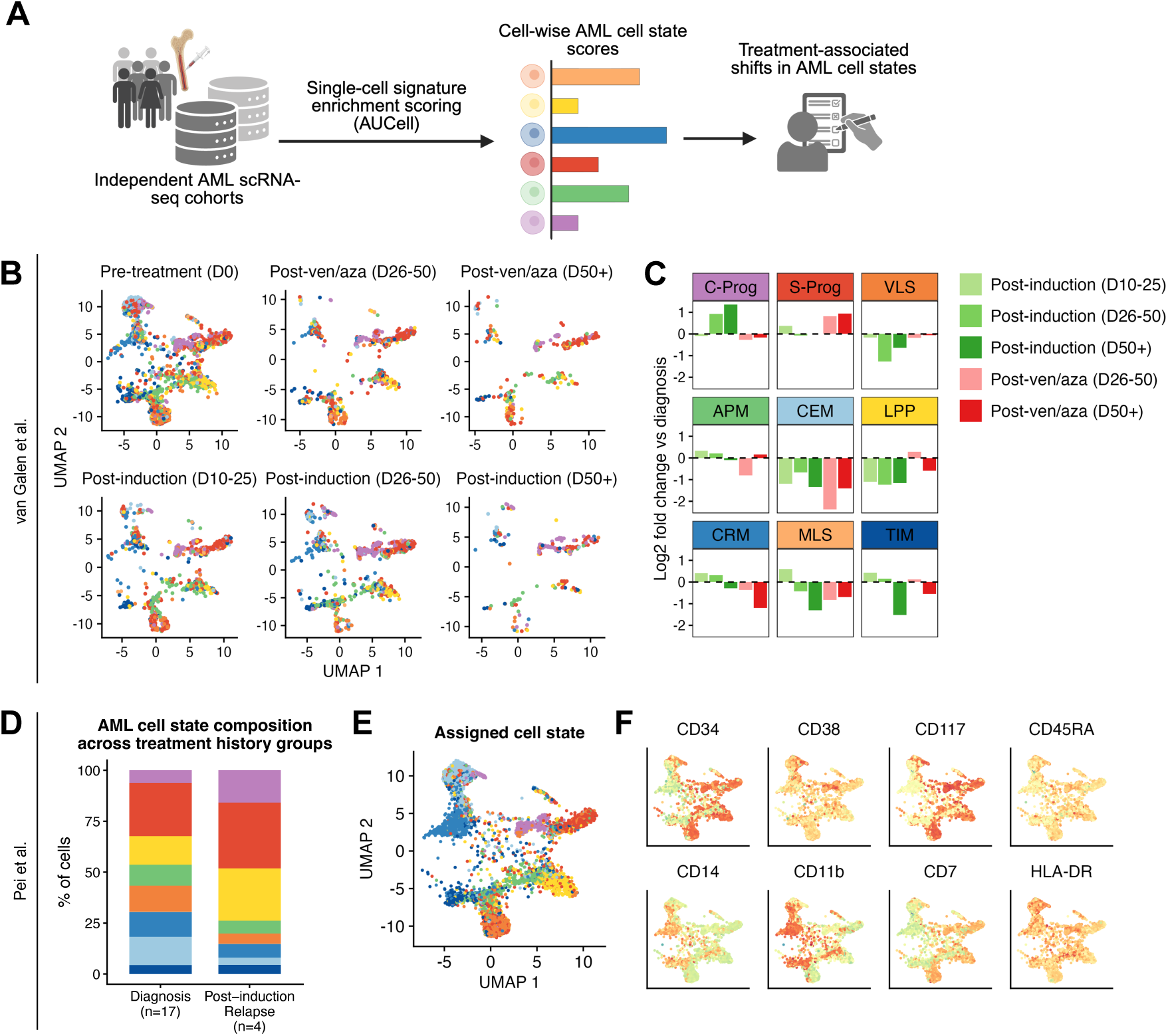
Selective persistence of AML cell states across treatment timepoints. (A) Visual representation of the analysis workflow. (B) Projected UMAPs showing cells in the van Galen dataset projected onto the original 20-gene signature UMAP coordinates, split by treatment type and timepoint, and where each cell is colored by its assigned cell state. (C) Bar plots showing log2 fold change in assigned cell states compared to diagnosis. (D) Bar plots showing the average proportion of cells (%) per treatment group. (E) Projected UMAP showing cells in the Pei dataset projected onto the original 20-gene signature UMAP coordinates, each cell is colored by its assigned cell state. (F) Projected UMAP overlaid with protein expression measured by CITE-seq.

In the dataset from van Galen et al., consisting of samples from diagnosis (n=16), post-induction chemotherapy (n=10), and venetoclax/azacitidine (ven/aza) treatment (n=1), projection onto our original UMAP coordinates revealed treatment-specific shifts in cell state composition (Figure 5B-C). Induction chemotherapy reduced LPP, CEM, and VLS. In contrast, C-Prog showed an increase with each post-induction timepoint. MLS and TIM showed an initial transient increase at early post-induction timepoints (D10-25) followed by gradual reduction. S-Prog, APM, and CRM remained largely unchanged over treatment timepoints.

Ven/aza treatment showed a distinct pattern of state-selective effects. CEM, CRM, and MLS were reduced, while C-Prog, S-Prog and VLS were increased or maintained. LPP and TIM were initially unchanged and showed a reduction in later timepoints, while APM showed the inverse.

We then utilized the dataset from Pei et al. where patients were either sampled at diagnosis (n=17) or at post-induction relapse (n=4). Although patients at diagnosis showed diverse cell state compositions, relapse samples showed similar profiles (Supplementary Figure 5A-B). Overall, relapse samples showed a lower proportion of monocytic cell states, and were mainly comprised of C-Prog, S-Prog, and LPP (Figure 5D). The dataset contained surface protein expression data partially overlapping with our CITE-seq panel (Figure 5E-F). Protein expression patterns across cell states were consistent with our previous observations, such as expected CD34 and CD14 distributions, and had increased CD38 and CD45RA expression in C-Prog assigned cells. CD7 was additionally increased in C-Prog, S-Prog, LPP, and VLS.

### Distinct AML cell states exhibit divergent microenvironmental signaling programs

Having established that AML cell states associate with drug sensitivity profiles and soluble protein signatures, we next explored the cell-cell communication programs that may underlie these functional differences. Using CellChat on our scRNA-seq data, we inferred outgoing and incoming signaling pathway activity and interaction probabilities between all nine AML cell states and the lymphocyte (T/B cell) population.

The monocytic states CEM, CRM, and TIM showed the highest number of interactions and interaction strength overall (Figure 6A-B, Supplementary Figure 6A), consistent with enrichment in inflammatory chemokine-associated plasma proteins (Figure 3D). Both incoming and outgoing interaction strengths were high for these states, suggesting they function as active signaling hubs within the microenvironment. Among progenitor states, S-Prog showed the highest outgoing interaction strength despite intermediate incoming strength.

**Figure 6:**
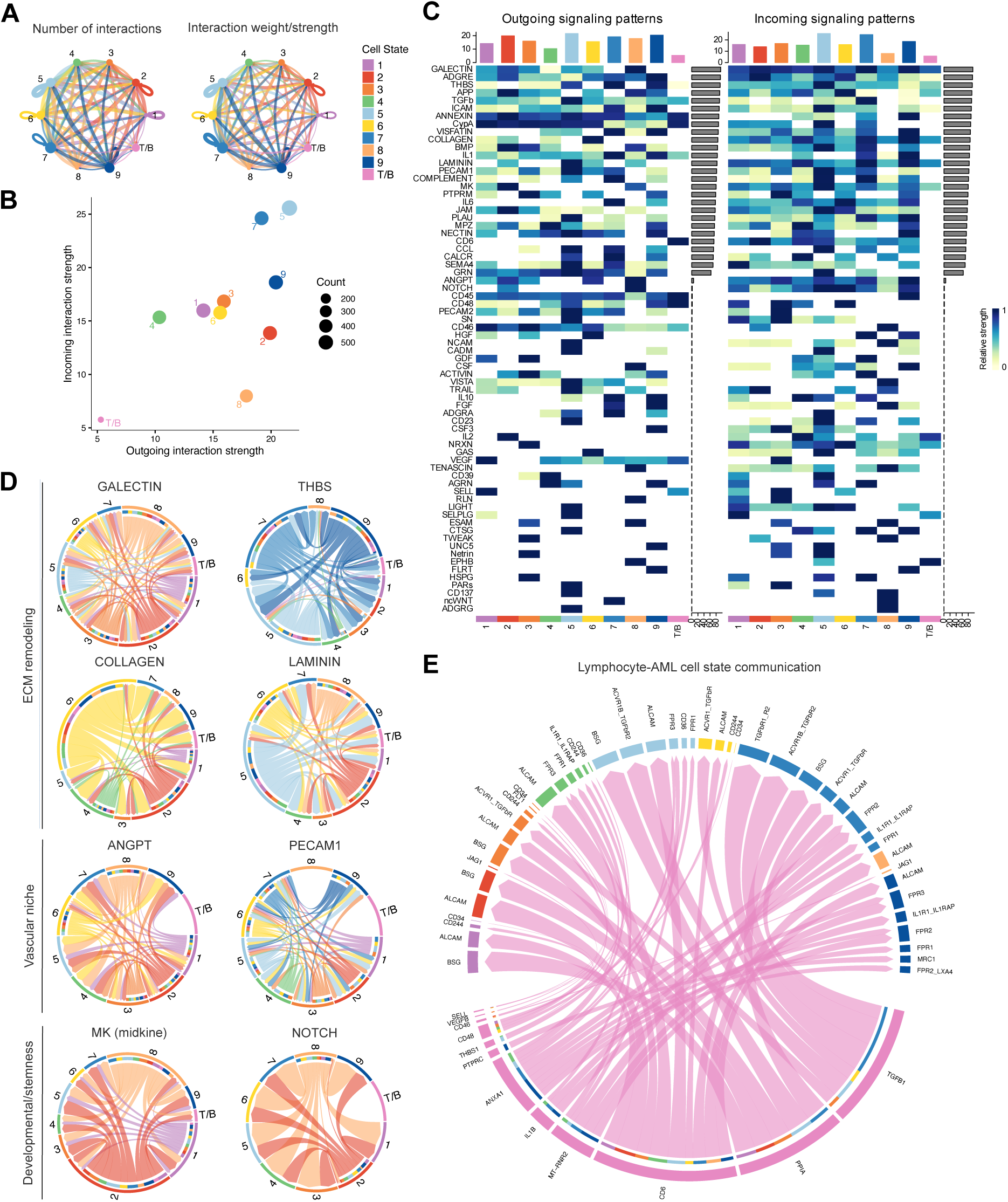
AML cell states exhibit divergent microenvironmental signaling programs. (A) The number of inferred interactions and their strengths for each cell state measured with CellChat. (B) XY plot of incoming and outgoing interaction strength for each cell state, with dot size representing gene count. (C) Heatmap with all outgoing (left) and incoming (right) signaling patterns showing all significant pathways. Values are shown as relative signaling strength, and bar plots on the Y-axis represent gene count per pathway. (D) Chord plots of selected significant pathways. (E) Chord plot of T/B cell signaling with all cell states.

Analysis of specific signaling pathways revealed state-specific preferences for ECM remodeling and vascular niche programs (Figure 6C-D). GALECTIN signaling was broadly active across most states, while monocytic states preferentially utilized THBS signaling for ECM remodeling. CEM additionally showed high LAMININ signaling alongside LPP and the stromal states. Progenitor states including LPP, APM, and S-Prog showed prominent COLLAGEN pathway activity, while vascular niche signaling through ANGPT was restricted to progenitor-like states, with monocytic and APM states preferentially utilizing PECAM1.

MK and NOTCH signaling shared between S-Prog and MLS suggests a convergent stemness maintenance program between these transcriptionally distinct states. VLS showed a progenitor-like interaction profile despite its stromal transcriptional identity, with active ANGPT and PECAM1 vascular niche signaling.

Finally, analysis of lymphocyte interactions revealed that T/B cell communication was predominantly mediated by monocytic states (Figure 6E), consistent with the depletion of immune activation markers in CEM-enriched patients observed in our soluble proteomics data. Progenitor states with the exception of S-Prog showed shared CD244 and CD34-mediated interactions with lymphocytes, while S-Prog and MLS shared JAG1-mediated T/B cell communication, a Notch ligand interaction consistent with their shared inferred NOTCH pathway activity.

### Ligand-receptor analysis reveals state-specific niche interactions

To complement the pathway-level communication analysis performed with CellChat, we applied CellPhoneDB to our scRNA-seq data to detect possible specific ligand-receptor interactions between AML cell states. Analysis of interaction counts and strengths confirmed the broadly active signaling role of monocytic states identified earlier, with CEM, CRM, and TIM showing the highest number of significant interactions across all cell state pairs (Supplementary Figure 7A-B). Consistent with their associated enrichment in inflammatory chemokine-associated plasma proteins in our soluble proteomics analysis, monocytic states were identified as major signaling hubs, both sending and receiving signals across the AML cell state network.

We examined possible interactions between monocytic states and lymphocytes to investigate the immunosuppressive factor associations indicated by our soluble proteomics analysis, where CEM-enriched patients showed depletion of immune activation markers and elevated HO-1. CellPhoneDB analysis identified consistent interactions between all three monocytic states and T/B cells, with NECTIN2-TIGIT and PVR-TIGIT (Figure 7A). TIGIT is an established inhibitory immune checkpoint receptor expressed on T and NK cells, and its ligands NECTIN2 and PVR are known to suppress cytotoxic immune responses when expressed on tumor cells.^26^ CD86-CD28 and ICOSLG-ICOS costimulatory interactions were also identified across all three states, as well as TGFB1-TGFBR3, consistent with TGFβ-mediated immune suppression. To investigate potential mechanisms underlying the broad kinase inhibitor resistance of the vascular-like stromal state identified in our drug sensitivity analysis, we examined ligand-receptor interactions received by VLS (Figure 7B). VLS received significant signals from multiple cell states through several pathways. Among the most notable were IL6-IL6 receptor interactions originating specifically from CRM. Additional survival-relevant interactions included CD44-TYROBP from CRM and TIM, and TGFB1-TGFbeta receptor1 from LPP. The APP-CD74 and PPIA-BSG interactions were broadly present across all sender states, suggesting signaling independent of specific state identity.

**Figure 7:**
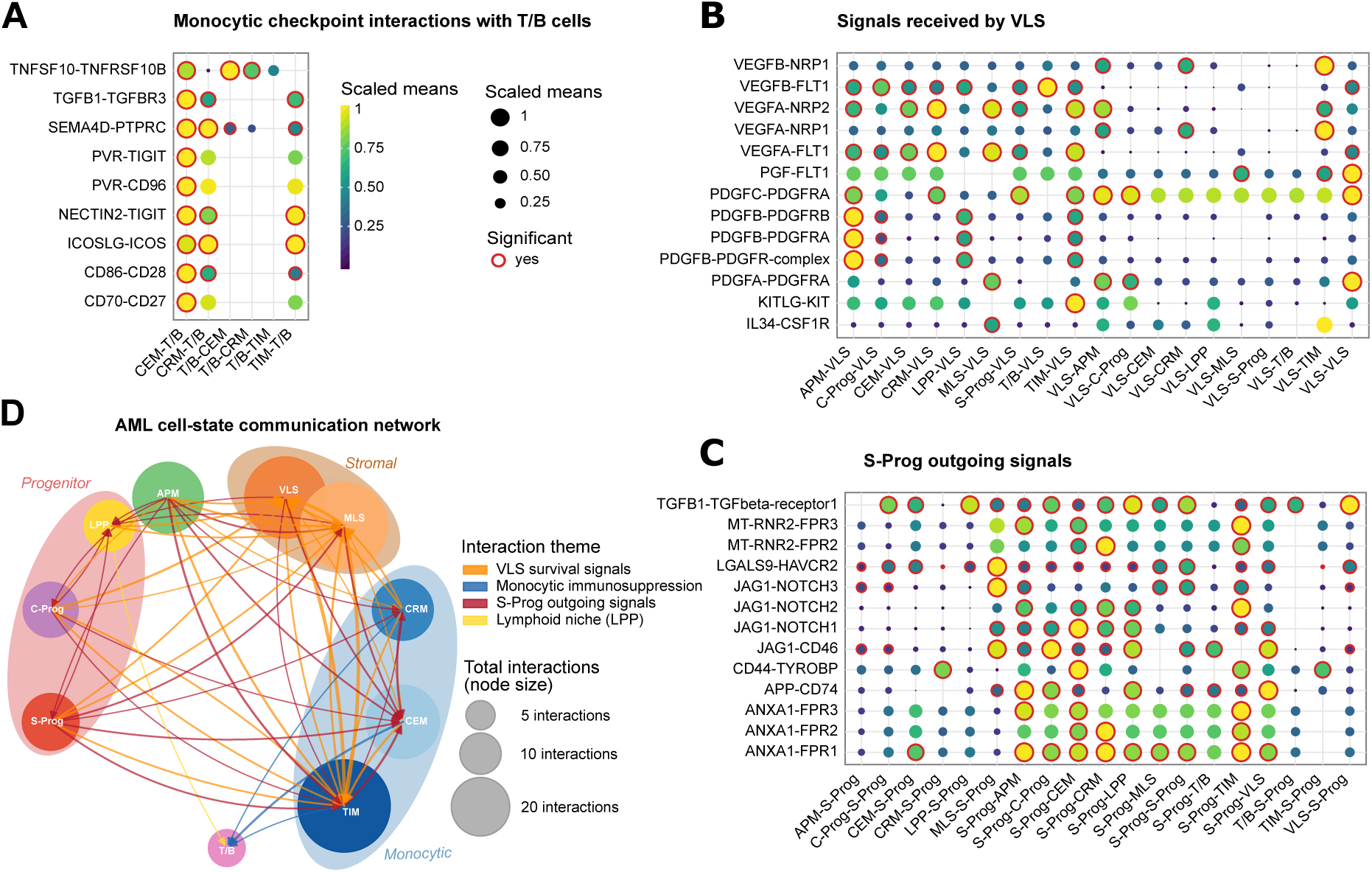
Cell-cell communication network between AML cell states. (A) Dot plot showing selected top cell-cell interactions between monocytic cell states and lymphocytes (T/B), (B) VLS and all other states, and (C) S-Prog with all other states. Color and size correspond to scaled mean expression while significance (p<0.05) is indicated by a red outline. (D) Directed network showing significant ligand-receptor interactions between AML cell states and T/B lymphocytes inferred by CellPhoneDB. Each node represents a cell state, colored by lineage identity. Node size reflects the total number of significant interactions (weighted degree). Directed edges indicate the sender-receiver directionality of significant interactions, with edge width proportional to the number of significant ligand-receptor pairs between each state pair. Edge color indicates the interaction theme. Only state pairs with three or more significant interactions are shown. Significance was defined as p<0.05 based on CellPhoneDB statistical analysis with 1,000 permutations.

Analysis of S-Prog outgoing interactions revealed a broad and diverse signaling program consistent with its high outgoing interaction strength identified by CellChat. S-Prog sent significant signals to most cell states through multiple pathways, including Notch, TGF-beta, and Galectin-related, as well as selective interactions of MT-RNR2 (Humanin) with FPR2 and FPR3, specifically to monocytic states (APM, CEM, CRM, TIM) but not to progenitor or stromal states (Figure 7C). Together these findings position S-Prog as a central signaling node. (Figure 7D).

## Discussion

In this study, we defined nine transcriptionally and functionally distinct AML cell states from single-cell RNA sequencing data and demonstrated their clinical and biological relevance across multiple independent analytical layers. By integrating single-cell transcriptomics with immunophenotypic profiling, extending to bulk RNA-seq-based clinical associations, ex vivo drug sensitivity, soluble proteomics, longitudinal single-cell datasets, and cell-cell communication analysis, we provide a classification of functional AML heterogeneity that extends beyond the classical leukemic stem cell lineage paradigm. Emerging evidence has challenged this classical LSC definition, with multiple studies demonstrating that leukemia-initiating capacity, therapy resistance, and transcriptional stemness programs are not confined to a single immunophenotypic compartment.^8,9,27^ Our findings extend this challenge by demonstrating that diagnostic stemness-associated transcriptional programs, including LSC17, APS, and iScore, are distributed across multiple cell states rather than a single population. The the cell states that we define, including stromal-like (VLS, MLS) and monocytic-specialized (CEM, CRM, TIM) programs alongside progenitor states (LPP, S-Prog, C-Prog) as well as unique mixed phenotype populations (APM), is consistent with emerging single-cell atlas studies showing that AML recapitulates a broad spectrum of myeloid differentiation programs within individual patients.^18,28^ This study build on previous studies^17–21,29^ which characterized the transcriptional architecture of AML at single-cell resolution, by providing functional annotations of these states through multi-dimensional clinical and pharmacological validation rather than transcriptional characterization alone.

LPP and S-Prog emerged as the most consistently stem-like states across independent scoring frameworks, yet their clinical associations were different. LPP enrichment characterized lower disease burden and specific molecular contexts including RUNX1 mutation, while S-Prog enrichment associated with adverse clinical features, older age, and a trend toward inferior survival. These opposing clinical profiles despite shared stemness signatures suggest that transcriptional stemness is a heterogeneous property that manifests differently depending on the broader cell state context.

A central contribution of this work is that transcriptional cell state enrichment associate with ex vivo drug sensitivity. The convergent BH3 mimetic resistance of all three monocytic states (CEM, CRM, and TIM) is consistent with prior reports describing venetoclax resistance in monocytic AML, including the identification of a distinct monocytic LSC population with reduced BCL2 dependence by Pei et al.^20,28^ The identification of S-Prog sensitivity to Wee1 and PLK1 inhibitors represents therapeutic candidates in a state associated with treatment resistance and adverse prognosis. The stress-adapted transcriptional identity of S-Prog may render these cells dependent on cell cycle checkpoint mechanisms to manage replicative stress, a dependency that could be exploited. Similarly, the association between C-Prog enrichment and Crenolanib sensitivity, which was observed largely independently of FLT3-ITD status, raises the possibility that cycling progenitor state enrichment could serve as a functional biomarker for kinase/FLT3 inhibitor sensitivity beyond mutation-based stratification alone, consistent with clinical trials of FLT3 inhibitors in FLT3-WT.^30^

Cell-cell communication analyses along with plasma proteomics data indicate that AML cell states are not passive participants in the bone marrow microenvironment but may actively engage in complex bidirectional signaling programs. The identification of monocytic states as major immunosuppressive hubs, engaging T/B cells through NECTIN2/PVR-TIGIT checkpoint interactions, TGFβ signaling, and CD86/CD28 costimulatory pathways, provides a hypothesis for the depletion of immune activation markers observed in CEM-enriched patients in our plasma proteomics analysis. TIGIT is an established immune checkpoint increasingly recognized as clinically relevant in AML, and our findings suggest that monocytic cell state enrichment may identify patients most likely to benefit from TIGIT-targeted immunotherapy.^31,32^ This hypothesis is supported by the convergent immunosuppressive signaling across all three monocytic states, suggesting that targeting TIGIT ligands expressed on monocytic AML cells could broadly address immune evasion regardless of the specific monocytic program present. The broad kinase inhibitor resistance of VLS in combination with our cell-cell communication findings is intriguing. VLS received significant survival signals from monocytic states through IL-6/IL-6 receptor, TGFB1, CD44/TYROBP, and PLAUR/integrin pathways, suggesting that its resistance to targeted kinase inhibitors may be partly sustained by niche-derived cytokine signaling rather than being exclusively cell-intrinsic. This model, in which microenvironmental crosstalk sustains resistance to targeted therapy, is well established in solid tumors and chronic myeloid leukemia but has been less systematically characterized in AML at the cell state level.^33,34^ The divergent microenvironmental interaction profiles of VLS and MLS despite their shared stromal identity is noteworthy. VLS showed a progenitor-like communication pattern with active vascular niche signaling through ANGPT and PECAM1 pathways, while MLS showed higher outgoing interaction strength with shared stemness signaling through MK and NOTCH with S-Prog. These differences, along with differential drug response profiles and clinical associations, suggest that the two stromal-like states occupy distinct functional niches within AML where VLS is involved in vascular niche interactions and MLS engages in stemness-associated crosstalk.

The identification of S-Prog as a major signaling hub with selective MT-RNR2/Humanin cytoprotective signaling to monocytic states represents a hypothesis in which the stress-adapted progenitor state may support the survival of drug-resistant populations in the AML microenvironment. Humanin is a mitochondria-derived peptide with established cytoprotective and anti-apoptotic activity that signals through formyl peptide receptors FPR2 and FPR3, which are preferentially expressed on monocytic cells.^35–37^ While Humanin signaling has been described in aging-related diseases, its role in AML microenvironmental crosstalk has not been previously characterized to our knowledge.^38^ This warrants direct experimental validation.

Beyond individual signaling interactions, our data collectively support a model in which distinct AML cell states occupy specialized niche positions within the bone marrow microenvironment. VLS, with its vascular-like transcriptional identity and active ANGPT/PECAM1 vascular niche signaling, appears to occupy an endothelial niche position where it receives survival signals from surrounding monocytic and stromal states that may sustain its kinase inhibitor resistance independently of cell-intrinsic mechanisms.^39,40^ S-Prog, with its high outgoing interaction strength, stress-adapted identity, and JAG1-mediated Notch signaling to all surrounding states, resembles a niche organizer and may be actively remodeling its microenvironment in ways that may create permissive conditions for its own persistence and that of drug-resistant monocytic states.^41^ MLS, sharing stemness-associated MK and NOTCH signaling programs with S-Prog despite its stromal transcriptional identity, may represent a bone marrow mesenchymal stromal cell-like program that supports leukemic cell maintenance.^42^

## Methods

### Primary AML sample processing

BM or PB from AML patients was collected after obtaining informed written consent at the Karolinska University hospital and the Acute Leukemia Biobank. This study was approved by the regional Ethical Review Authority in Stockholm (DNR: 2017/2085-31/2, 2025-08490-01) and adhered to the Declaration of Helsinki. Sample processing and mononuclear cell isolations were performed as described previously.^10,43^ Supernatants for soluble proteomic analysis were obtained by centrifugation of diluted aspirates in RPMI at 300 g for 10 minutes and stored at –80°C until analysis.

### scRNAseq and CITE-seq

Single-cell RNA sequencing with paired surface protein profiling (CITE-seq) was performed on primary AML samples using the 10x Genomics Chromium platform. Libraries included gene expression (GEX), antibody-derived tags (ADT), and hashtag oligonucleotides (HTO) used for sample multiplexing. Raw sequencing data were processed using the Cell Ranger multi pipeline (version 9.0.1, 10x Genomics), which performs joint alignment and quantification of gene expression and feature barcode libraries.

Following initial processing with Cell Ranger, HTO counts were extracted and demultiplexing was performed using the HashSolo algorithm as implemented in Scanpy. HashSolo applies a probabilistic model to assign each cell to a sample of origin based on HTO signal, while simultaneously identifying doublets and negative droplets. Only cells classified as high-confidence singlets were retained for downstream analysis. Gene expression quantification was based on unique molecular identifiers (UMIs) generated by the Cell Ranger pipeline. Cells were filtered based on standard quality control metrics to remove low-quality or damaged cells. Specifically, cells were only kept if they had at least 3000 total UMIs, at least 1500 detected genes and less than 15% mitochondrial reads. Cells failing any of these criteria were excluded from downstream analysis. In addition, genes were required to be detected in at least 10 cells to be retained for downstream analyses. CD3, CD16, CD4, CD11c, CD56, CD14, CD8, CD45, CD117, CD25, CD123, CD34, CD15, and HLA-DR.

### Single-cell profiling and cell type annotation

Single-cell RNA-seq data were processed using Seurat (v5.3.1), including log-normalization (scale factor 10,000), identification of 2,000 variable features, PCA, UMAP, and graph-based clustering (resolution 0.5). Cell type annotation was performed by mapping query cells to the BoneMarrowMap^21^ (v0.1.0) reference using Symphony (v0.1.2), with patient ID used as a batch covariate. Cells with high mapping error (MAD > 2.5) were excluded. Gene set activity scores were computed using AddModuleScore with curated AML and hematopoietic gene signatures. Cluster marker genes were identified using the Wilcoxon rank-sum test (min.pct = 0.25, logFC > 0.25), and higher-order structure was defined by k-means clustering on principal components, with k selected based on silhouette analysis. Normalized antibody-derived tag (ADT) data from CITE-seq were matched to scRNA-seq cells based on shared cellular barcodes.

### Cell state gene signature generation

Marker genes for each meta-cluster were identified using the Seurat FindAllMarkers function (Wilcoxon rank-sum test; min.pct = 0.25, logFC > 0.25). Pathway enrichment analysis was performed using over-representation analysis (ORA) from clusterProfiler (v4.14.6), using MSigDB Hallmark gene sets. P-values were adjusted using the Benjamini-Hochberg method, and significantly enriched pathways (adjusted p < 0.05) were retained. To define meta-cluster-specific gene signatures, the top-ranked marker genes per cluster were selected and used for downstream analyses. These signatures were used to generate a reduced-dimensional representation (PCA and UMAP) and to assess cell state structure. Gene signature activity was quantified at the single-cell level using AUCell (v1.28.0). Cells were assigned to meta-clusters based on the highest AUCell score exceeding a predefined threshold. To assess the specificity of the gene signatures, AUCell-based classification accuracy was compared to randomly generated gene sets of matched size across multiple iterations.

### Bulk RNAseq

Sample preparation and sequencing were performed as previously described.^44^ RNA-seq data were processed using a reproducible workflow implemented in Snakemake, available at https://github.com/lhartmanis/RNA-seq_processing_pipeline. Raw paired-end FASTQ files were subjected to quality control using FastQC^45^, and adapter sequences and low-quality bases were removed using fastp.^46^

Reads were aligned to the human reference genome (GRCh38) using the splice-aware aligner STAR^47^ (v2.7.3a), with gene annotation from GENCODE v45. Alignment was performed in paired-end mode with sjdbOverhang=99 and outFilterMultimapNmax=50, and gene-level counts were generated using the ––quantMode GeneCounts option. Gene expression was quantified using featureCounts^48^ with the parameters –p ––largestOverlap ––primary, assigning reads to both exonic and intronic regions to maximize coverage and sensitivity.

### Bulk RNA-seq signature scoring

Bulk RNA-seq data were obtained from an integrated cohort of AML samples with prior batch correction performed using ComBat following removal of mitochondrial and ribosomal genes. Gene expression counts were filtered using edgeR (v4.4.2), normalized using trimmed mean of M-values (TMM), and transformed to log2 counts per million (logCPM). Meta-cluster gene signatures derived from single-cell analyses were scored in bulk samples using gene set variation analysis (GSVA) (v2.0.7). Only AML samples with available clinical annotation were retained for downstream analyses.

### Clinical associations

Clinical data was obtained through patient records and from the Swedish Adult Acute Leukemia Registry. Associations between signature scores and clinical variables were assessed using non-parametric statistical tests using rstatix (v0.7.3), survival (v3.8-3), and survminer (v0.5.1). Spearman correlation was used for continuous variables, Wilcoxon rank-sum tests for binary variables, and Kruskal-Wallis tests for multi-category variables. Where appropriate, post hoc pairwise comparisons were performed using Dunn’s test. P-values were adjusted for multiple testing using the Benjamini-Hochberg method. Overall survival analysis was performed using Kaplan-Meier estimation with log-rank testing. Patients were stratified into high and low groups based on median signature scores.

### Drug sensitivity and resistance testing

DSRT was performed as described previously.^43^ In short, cells were resuspended in RPMI medium (RPMI 1640, 10% FBS, 2mM L-glutamine, 100 IU/mL Penicillin, 0.1 mg/mL Streptomycin) and dispensed in 384-well drug plates containing 528 oncological compounds (FIMM HTB). Incubation was done at 37°C and 5% CO_2_ for 72h, and cell viability was evaluated with CellTiterGlo (CTG, Promega, Madison, WI). Luminescence was quantified with an EnSight plate reader (PerkinElmer) and drug sensitivity scores (DSS) were calculated with Breeze.^49^

### Drug sensitivity associations

Spearman correlation was used to assess associations between GSVA-derived cell state enrichment scores and drug response scores (sDSS) across patients. Drugs tested in fewer than 76 samples were excluded. P-values were adjusted for multiple testing using the Benjamini-Hochberg method. For selected drugs of interest, patients were stratified into high versus low cell state groups (median split), and differences in drug sensitivity were assessed using the Wilcoxon rank-sum test.

### Olink soluble proteomics

Protein abundance profiling was performed using a Proximity Extension Assay (PEA) based on the Olink 96 Immuno-Oncology panel (Olink Proteomics, Uppsala, Sweden), following the manufacturer’s protocol. Protein quantification was carried out on the BioMark HD system (Fluidigm), and raw Ct values were processed using Fluidigm Real-Time PCR Analysis software. Normalization and quality control were performed using the Olink NPX Manager pipeline, incorporating internal and external controls to generate log2-scaled Normalized Protein eXpression (NPX) values.

### Plasma proteomics (Olink) associations

Spearman correlation was used to assess associations between GSVA-derived AML cell state scores and Olink plasma protein expression data. Only samples present in both datasets were included in the analysis. Multiple testing correction was performed using the Benjamini-Hochberg method.

### Single-cell mapping of AML cell states across external patient cohorts

Single-cell RNA-seq datasets (van Galen et al.^17^ and Pei et al.^20^) were analysed using Seurat. Cells were filtered for low-quality clusters and processed by normalization, variable feature selection, PCA, and UMAP embedding, followed by graph-based clustering. For the Pei dataset, CITE-seq ADT data were integrated and used for marker validation. AML cell states were defined using previously curated gene signatures, and per-cell state activity scores were computed using AUCell based on ranked gene expression. Cell state labels were assigned by maximum signature enrichment. The datasets were additionally mapped to a the BMM atlas using Seurat label transfer to infer cell type identities, followed by quality control based on mapping error scores.

### Inference of cell-cell communication networks

Cell-cell communication was inferred using CellChat (v.2.2.0) on quality-controlled AML single-cell RNA-seq datasets. Cells were grouped into annotated AML and immune/stromal compartments based on previously defined cell-state labels. Gene expression matrices were used to construct CellChat objects, and the human CellChatDB database was applied for ligand-receptor analysis. Overexpressed genes and interactions were identified, and communication probabilities were computed using the trimean approach followed by pathway-level aggregation. Global and pathway-specific interaction networks were inferred, and interaction strength was quantified based on aggregated edge weights. Network topology and interaction contributions were further assessed to identify dominant sender-receiver relationships across AML and microenvironmental compartments.

Cell-cell communication was further inferred using CellPhoneDB (v5.0.0) in Python (v3.8.20) within a Conda-managed Jupyter environment. Normalized gene expression matrices and annotated cell-type labels were used as input to construct ligand–receptor interaction networks. Statistical analysis was performed using permutation-based testing (1,000 iterations) to identify significant interactions between cell populations, applying a 10% expression threshold and p-value cutoff of 0.05. Interaction scores were computed across all annotated AML and microenvironmental compartments using the CellPhoneDB human ligand-receptor database. Significant interaction pairs and their mean expression values were aggregated to generate intercellular communication maps.

### Data analysis and visualization

Plots were generated with R (v4.4.3) using RStudio (v2024.12.1+563) with ggplot2 (v4.0.0), ggrepel (v0.9.6), cowplot (v1.2.0), patchwork (v1.3.2), and RColorBrewer (v1.1-3) unless stated otherwise. CellPhoneDB plots were generated in Python (v3.8.20) using Seaborn (v0.13.2) and Matplotlib (v3.7.5)

### Data Sharing

Drug testing, soluble proteomics, and single cell data sets will not available publicly due to privacy reasons. Access to the data at the Swedish National Data Service will be approved pending ethical compliance.

### Code Availability Statement

All analysis code can be found on GitHub at https://github.com/nstruyf/AML_cell_states. RNA-seq data processing can be found at https://github.com/lhartmanis/RNA-seq_processing_pipeline.

## Acknowledgements

We would like to thank all the participants in the study. We also thank Claudia Fredolini and Annika Bendes of the Affinity Proteomics unit at SciLifeLab, Stockholm, Sweden, for their support with the Olink data generation. This work was funded by the Swedish childhood cancer fund (TJ2021-0080) (T.E.), Karolinska Institutet (2024-00304, 2020-01091, 2018-02283) (T.E.), the Swedish foundation for strategic research (SB16-0058) (O.K.), the Swedish research council (2017-06095) (O.K.), and the Knut and Alice Wallenberg foundation (2015.0291) (O.K.).

## Authorship Contributions

Contribution: N.S. and T.E. conceptualized the study; N.S., L.R.P., and S.B. performed data generation; N.S., L.H., and A.Ö. performed data processing; N.S. performed data analysis and made figures; S.L., O.K., and T.E. contributed to the study supervision; A.B., S.B., S.L., O.K., and T.E. provided study resources; N.S. wrote and edited the original manuscript draft

## Conflict-of-interest disclosures

None of the authors have conflicts of interest to declare.

Correspondence: Nona Struyf; Department of Oncology-Pathology, Karolinska Institute, SciLifeLab, Tomtebodavägen 23, 17165, Solna, Sweden, Tom Erkers; Department of Oncology-Pathology, Karolinska Institute, SciLifeLab, Tomtebodavägen 23, 17165, Solna, Sweden;

## Notes

### Competing Interest Statement

The authors have declared no competing interest.

