## Supplemental material for "Transcriptionally defined AML cell states associate with treatment response and microenvironmental remodeling"

### Supplemental Figures

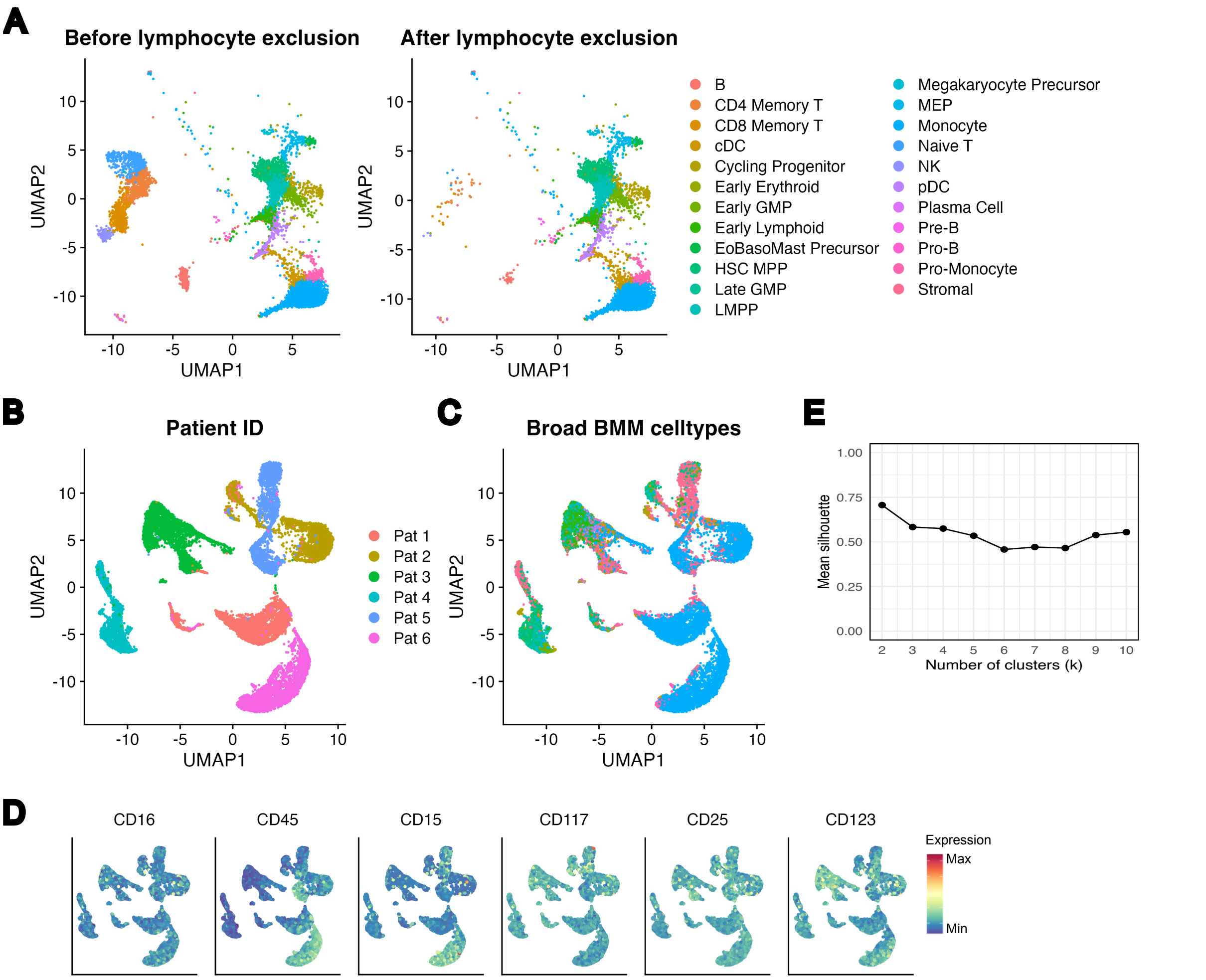


**Supplementary Figure 1:** (A) UMAPs showing cells projected on BMM coordinates before and after removal of lymphocytes. (B) UMAPs of primary clusters colored by patient ID and (C) Broad BMM celltype assignment. (D) UMAPs overlaid with surface protein expression measured by CITE-seq. (E) Silhouette plot for meta-clustering showing mean silhouette up to k=10.

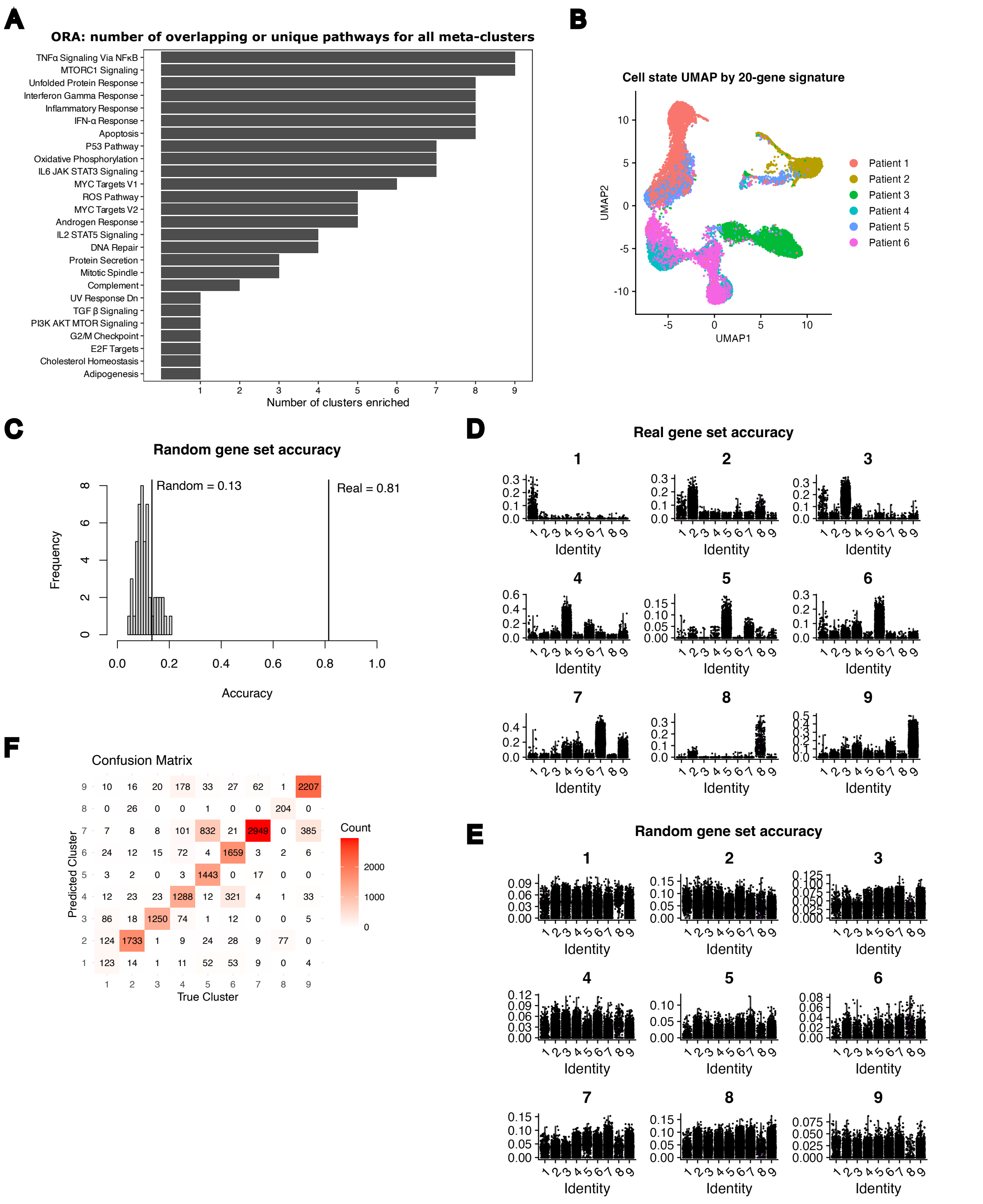


**Supplementary Figure 2:** (A) Plot showing the number of meta-clusters where enrichment occurred in the ORA. (B) UMAP clustered by 20-gene signature colored by patient. (C) Bar plot showing accuracy of a random selected gene set vs actual predicted accuracy. (D) Real and (E) random gene set accuracy shown per meta-cluster. (F) Confusion matrix showing predicted (rows) versus true (columns) cluster labels. Counts are indicated by color and numbers. Random gene signatures (size-matched, 50 iterations) were used as a control to estimate baseline classification accuracy.

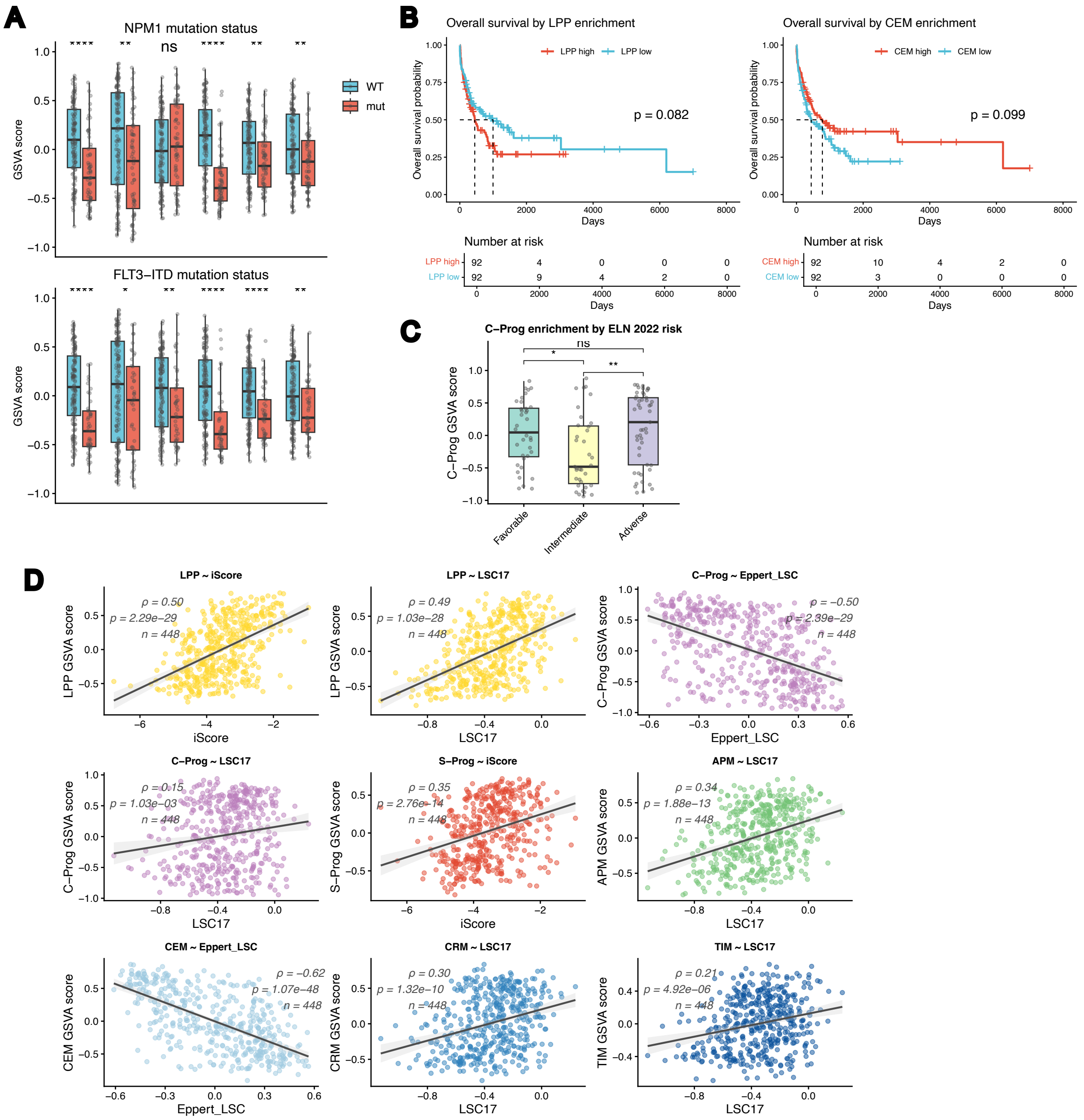


**Supplementary Figure 3:** (A) Bar plots with GSVA scores for all cell states compared to mutation status of NPM1 (top) and FLT3-ITD (bottom). (B) Kaplan-Meier curves for LPP and CEM. (C) C-Prog GSVA score compared to ELN 2022 status. (D) Dot plots showing associations of cell states to existing diagnostic or classical LSC/HSC signatures.

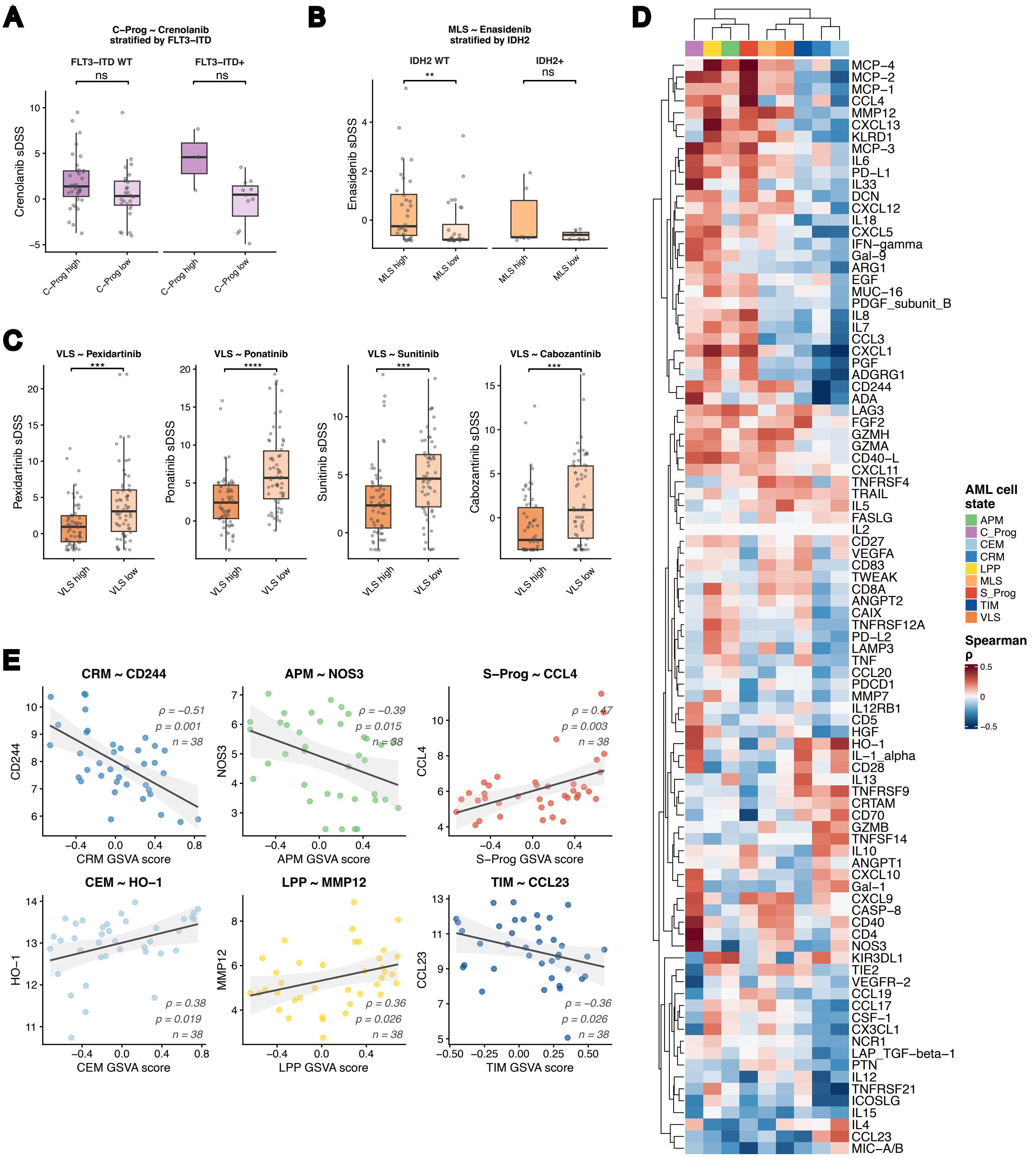


**Supplementary Figure 4:** (A-B) Boxplots of selected drug-cell state associations compared to mutational status, significance was assessed with the Wilcoxon test. (C) Boxplots of selected drug-VLS associations, significance was assessed with the Wilcoxon test. (D) Heatmap of all tested soluble proteins in association to cell state GSVA scores displayed as Spearman ρ. (E) Scatterplots showing association between selected soluble proteins and cell state GSVA scores for all tested patients (n=38), assessed with Spearman correlation.

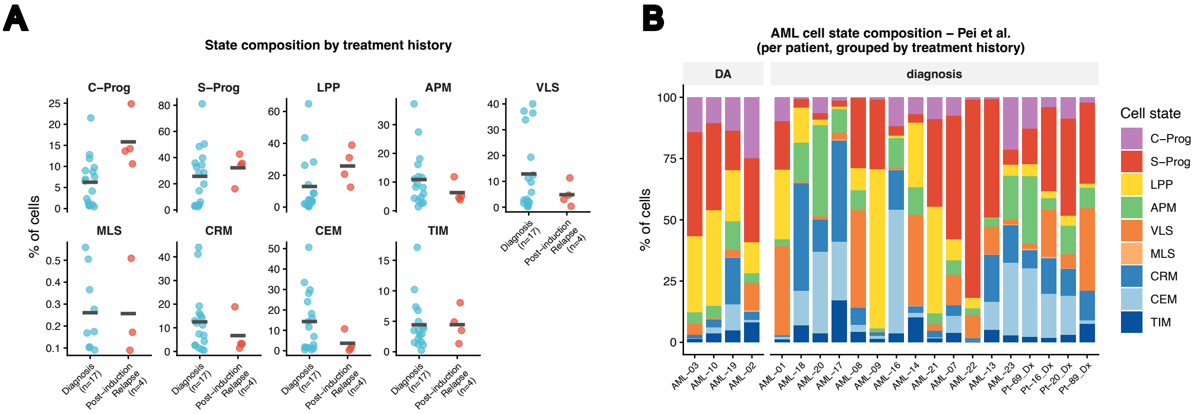


**Supplementary Figure 5:** (A) Dot plots and (B) bar plots showing the proportion of cells (%) for each patient per treatment group in the Pei et al. dataset, split by cell state.

A

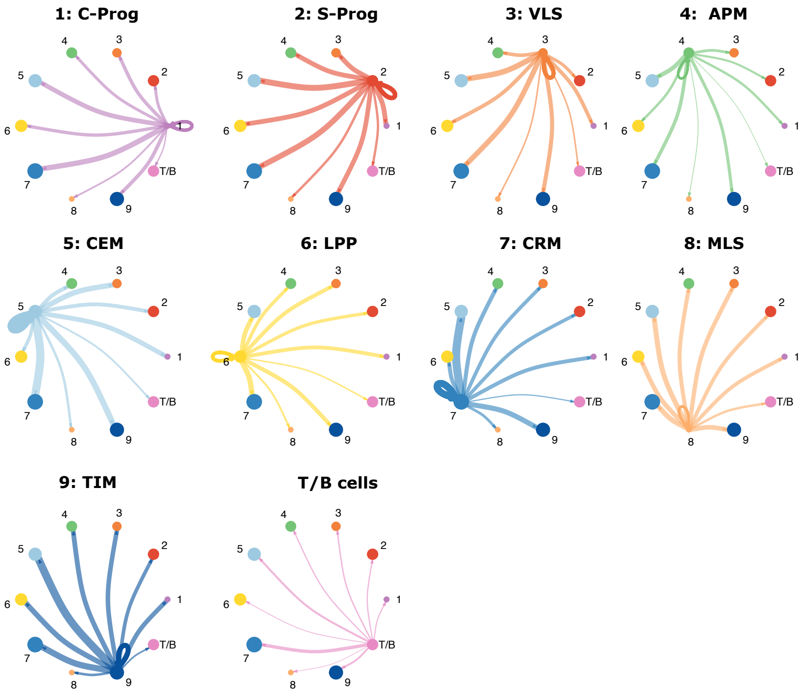


**Supplementary Figure 6:** (A) The number of inferred interactions and their strengths for each cell state measured with CellChat, split by cell state.

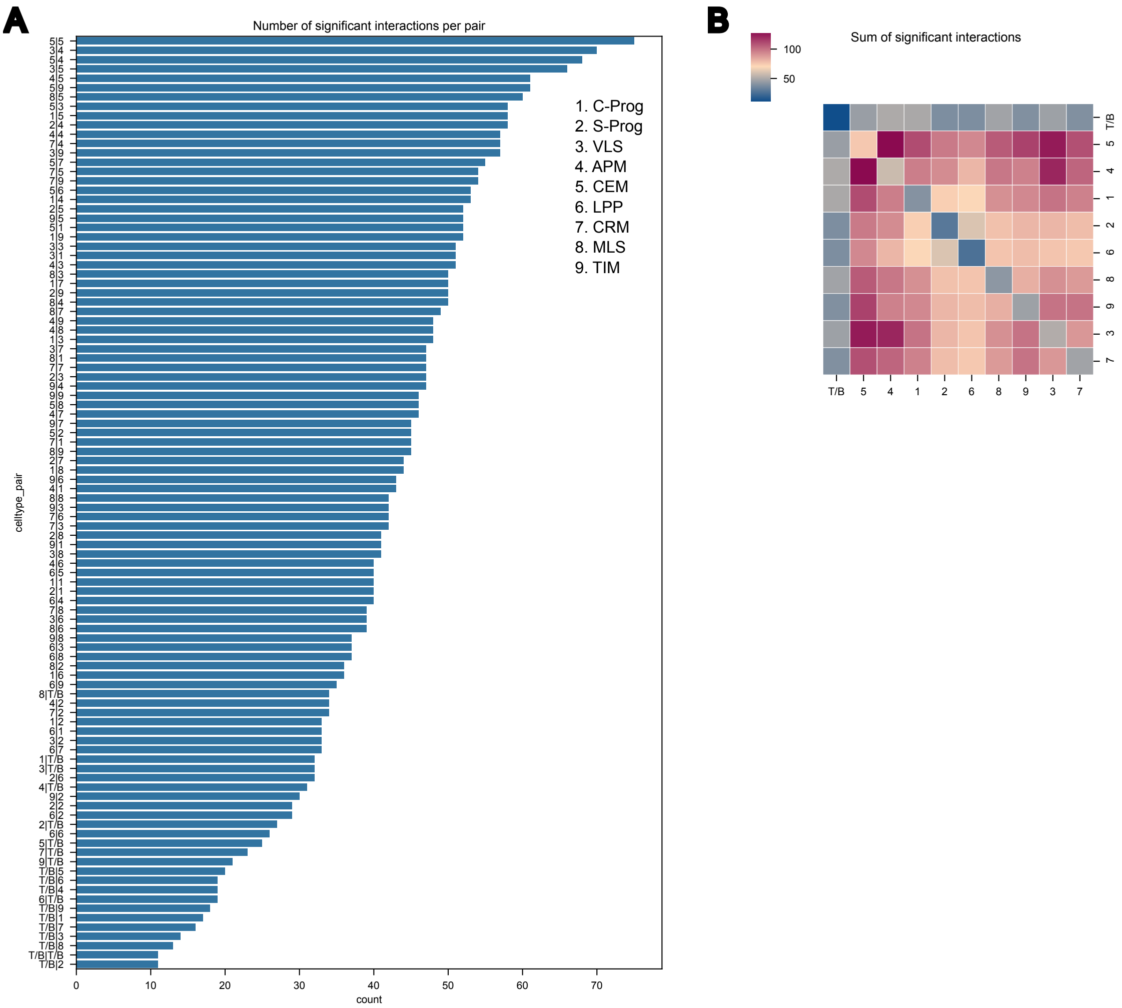


**Supplementary Figure 7:** (A) Bar plot and (B) heatmap showing the number of significant interactions per cell state pair according to CellPhoneDB.
